# SARS-CoV-2 Neutralizing Antibodies Following a Second BA.5 Bivalent Booster

**DOI:** 10.1101/2023.08.13.553148

**Authors:** Qian Wang, Anthony Bowen, Jerren Ho, Richard Zhang, Riccardo Valdez, Emily Stoneman, Aubree Gordon, Lihong Liu, David D. Ho

**Author notes:** Correspondence (A.G.), (L.L.), (D.D.H.). These authors contributed equally (Qian Wang, Anthony Bowen, Jerren Ho, Richard Zhang).

## Abstract

Bivalent COVID-19 mRNA vaccines expressing both the ancestral D614G and Omicron BA.5 spike proteins were introduced in August 2022 with the goal of broadening immunity to emerging SARS-CoV-2 Omicron subvariants. Subsequent studies on bivalent boosters found neutralizing antibody responses similar to boosters with the original monovalent vaccine, likely the result of immunological imprinting. Guidelines allow for administration of a second bivalent booster in high-risk groups, but it remains unknown whether this would broaden antibody responses. To address this question, we assessed longitudinal serum SARS-CoV-2-neutralizing titers in 18 elderly immunocompetent individuals (mean age 69) following a fourth monovalent booster and two BA.5 bivalent booster vaccines using pseudovirus neutralization assays against D614G, Omicron BA.5, and Omicron XBB.1.5. There was a small but significant increase in peak neutralizing antibody responses against Omicron BA.5 and XBB.1.5 following the first bivalent booster, but no significant increase in peak titers following the second bivalent booster. Omicron-specific neutralizing titers remained low after both doses of the BA.5 bivalent booster. Our results suggest that a second dose of the BA.5 bivalent booster is not sufficient to broaden antibody responses and to overcome immunological imprinting. A monovalent vaccine targeting only the spike of the recently dominant SARS-CoV-2 may mitigate the “back boosting” associated with the “original antigenic sin.”

## Main text

To enhance immunity to evolving severe acute respiratory coronavirus 2 (SARS-CoV-2) variants, updated bivalent mRNA vaccines targeting both the Omicron BA.5 and ancestral D614G spike proteins were introduced in August 2022. However, we and others found that serum neutralizing antibody (NAb) titers against SARS-CoV-2 variants were similar approximately one and three months following either bivalent or monovalent booster, underscoring the presence of immunological imprinting^1–4^. Guidelines now allow for a second BA.5 bivalent vaccine shot for immunocompromised and older adults, although whether an additional booster can overcome immunological imprinting remains unknown. This study was conducted to answer this question.

Here, we assessed serum SARS-CoV-2-neutralizing titers in 70 longitudinal samples from 18 immunocompetent individuals following multiple mRNA vaccine doses to track antibody dynamics over time. All participants received four monovalent mRNA vaccines followed by two bivalent boosters (**Table S1**). Sera were collected ∼1 month following the fourth monovalent (M4) vaccine, ∼1 and ∼6 months after the first bivalent (B1) booster, and ∼1 month after the second bivalent (B2) booster (**Figure 1A**). NAb titers (ID_50_) were determined against D614G, BA.5, and XBB.1.5 using a pseudovirus neutralization assay (see Supplementary Appendix). Participants had a mean age of 68.9 years and no known infection history (**Table S2**). One serum sample was excluded due to the presence of anti-nucleocapsid antibodies (**Figure S1**).

**Figure 1.**
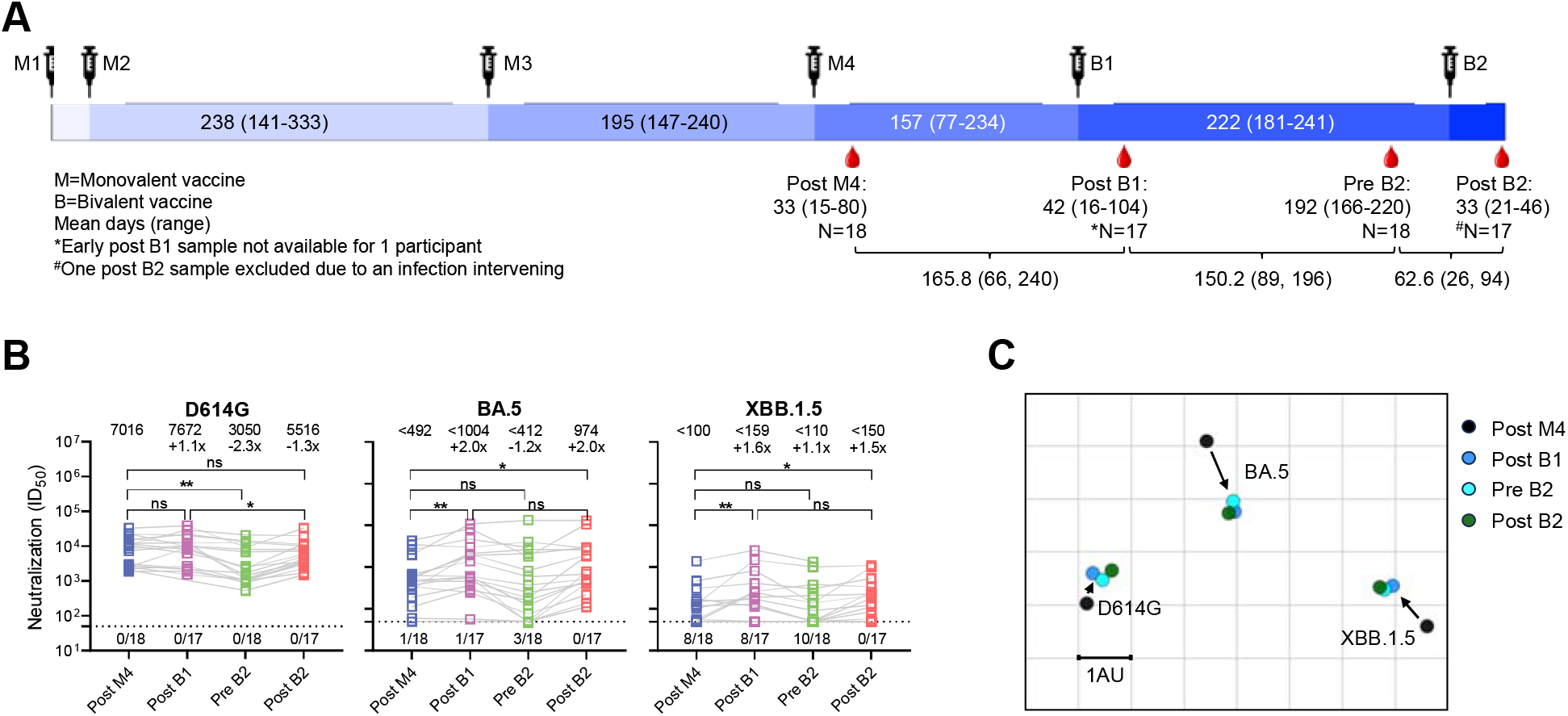
SARS-CoV-2 NAb responses following a second BA.5 bivalent booster. **(A)** Longitudinal samples were collected from 18 individuals ∼1 month following the fourth monovalent (M4) vaccine, ∼1 and ∼6 months following first bivalent (B1) booster, and ∼1 month following the second bivalent (B2) booster. Timepoints are illustrated with the mean and range for the cohort. **(B)** Neutralizing ID_50_ titers at each timepoint against D614G, BA.5, and XBB.1.5. The geometric mean ID_50_ titer (GMT) and fold change compared to the post-M4 timepoint are denoted above each serum timepoint. Comparisons were made by Mann-Whitney tests (\**p* < 0.05; \*\**p* < 0.01). The assay limit of detection (LOD) is 50 (dotted line) and the number of samples at or below the LOD is denoted above the x-axis. **(C)** Antigenic maps generated for each serum timepoint were aligned to show changes in antigenic distance with each vaccine. Each antigenic distance unit (AU) represents an approximately a 2-fold change in ID_50_ titer.

There was no significant difference in NAb titers against the ancestral D614G at ∼1 month following the M4, B1, or B2 vaccine shots (**Figure 1B**). A small, but significant, increase in NAb titers was seen against Omicron BA.5 and XBB.1.5 ∼1 month following the B1 booster, but this difference did not persist over time, consistent with previous findings^3,4^. At ∼1 month following the B2 booster, peak NAb titers were similar to levels following the B1 booster. Peak NAb titers against both Omicron subvariants remained low in this cohort following both B1 and B2 boosters. Antigenic cartography based on these results showed that the B1 booster shortened the antigenic distances between D614G, BA.5, and XBB.1.5, while no change was seen following the B2 booster (**Figure 1C**).

These data provide compelling evidence that a repeat dose of the BA.5 bivalent mRNA vaccine, as currently recommended for certain populations, is not sufficient to broaden neutralizing antibody responses and to overcome immunological imprinting in elderly but healthy individuals. It is likely that removal of the ancestral spike from future COVID-19 vaccines could mitigate the “back boosting” due to the “original antigenic sin”, as suggested by modestly better serum NAb breadth in individuals following BA.5 infection^3^. Overall, these findings support current plans to update COVID-19 vaccines using only the spike protein of the latest dominant Omicron subvariant ^5^.

## Notes

### Competing Interest Statement

D.D.H. co-founded TaiMed Biologics and RenBio, serves as a consultant for WuXi Biologics and Brii Biosciences, and is a board director at Vicarious Surgical. Aubree Gordon served as a member of the scientific advisory board for Janssen Pharmaceuticals. The remaining authors have declared no conflicts of interest 

## References

1. Wang, Q. et al. Antibody Response to Omicron BA.4-BA.5 Bivalent Booster. New Engl J Med 388, 567–569 (2023).

2. Collier, A. Y. et al. Immunogenicity of BA.5 Bivalent mRNA Vaccine Boosters. New Engl J Med 388, 565–567 (2023).

3. Wang, Q. et al. SARS-CoV-2 neutralising antibodies after bivalent versus monovalent booster. Lancet Infect Dis 23, 527–528 (2023).

4. Lasrado, N. et al. Waning Immunity Against XBB.1.5 Following Bivalent mRNA Boosters. bioRxiv 2023.01.22.525079 (2023) doi:10.1101/2023.01.22.525079.

5. Updated COVID-19 Vaccines for Use in the United States Beginning in Fall 2023 | FDA. https://www.fda.gov/vaccines-blood-biologics/updated-covid-19-vaccines-use-united-states-beginning-fall-2023.

