## Supplemental information for "SARS-CoV-2 Neutralizing Antibodies Following a Second BA.5 Bivalent Booster"

### Supplementary Appendix

#### Supplementary Methods

*Clinical cohorts*

The longitudinal sera were obtained from the University of Michigan as part of the Immunity-Associated with SARS-CoV-2 Study (IASO), a continuing cohort study that started in Ann Arbor, Michigan, in 2020[1]. All IASO participants provided written informed consent, and serum samples were collected in accordance with the protocol approved by the Institutional Review Board of the University of Michigan Medical School. IASO participants complete weekly symptom surveys and are tested for SARS-CoV-2 upon report of symptoms.

In this study, sera were collected from 18 individuals with no recorded SARS-CoV-2 infections who had received four monovalent vaccines (Moderna mRNA-1273, Pfizer BNT162b2, or AstraZeneca ChAdOx1-S), followed by two doses of the BA.5 bivalent vaccine (Moderna mRNA-1273.222 or Pfizer BA.5 bivalent vaccine). Sera were collected approximately 30 days following the fourth monovalent vaccine dose (post M4), 30 days following the first bivalent vaccine dose (post B1), 6 months following first bivalent vaccine dose (pre B2), and 30 days following the second bivalent vaccine dose (post B2). Samples were examined by anti-nucleoprotein (NP) ELISA to confirm status of prior SARS-CoV-2 infection, with one positive sample excluded from analysis.

Clinical information for the study cohort is summarized in **Table S1** with detailed information for each individual in **Table S2**.

*Enzyme-linked immunosorbent assay (ELISA)*

The nucleoprotein of SARS-CoV-2 and the S2P spike trimer protein for D614G, BA.5, and XBB.1.5 were produced following methods that have been detailed previously[2].

To assess the binding levels of serum samples to the nucleoprotein and spike proteins, 50 ng/well of each protein was immobilized to ELISA plates. The plates were incubated them overnight at 4°C. Then, the ELISA plates were blocked with 300 µl of blocking buffer (1% BSA and 20% bovine calf serum (BCS) from Sigma) in PBS at 37°C for 2 hours. Then, the serum samples were gradually diluted by 5 times, starting with an initial 100× dilution, using a dilution buffer composed of 1% BSA and 20% BCS in PBS, and incubated on the ELISA plates at 37°C for 1 hour. For the anti-nucleoprotein (NP) ELISA test, serum samples from eight healthy donors collected pre-COVID pandemic were used as controls (referred to as "HD"). Following this, a 10,000-fold diluted Peroxidase AffiniPure goat anti-human IgG (H+L) antibody (from Jackson ImmunoResearch) was added and incubated for another hour at 37°C. The plates were washed between each step with PBST (0.5% Tween-20 in PBS). Lastly, the TMB substrate from Sigma was added and allowed to incubate until the reaction was stopped using 1 M sulfuric acid. The absorbance was then assessed at 450 nm, and $EC_{50}$ values were determined as the dilutions where the OD450 readings reached half of their maximum value, analyzed using GraphPad Prism version 9.2.

*Cell lines*

Vero-E6 cells (CRL-1586) and HEK293T cells (CRL-3216) were purchased from the American Type Culture Collection. Cells were grown in Dulbecco's Modified Eagle Medium (DMEM) containing 10% fetal bovine serum and 1% penicillin–streptomycin, in a humidified 5% CO2 atmosphere, at 37 °C.

*SARS-CoV-2 spike plasmids*

Plasmids that encode the spike (S) protein of SARS-CoV-2 variants D614G, BA.5, and XBB.1.5 have been previously constructed[3]. Before experimental use, the sequence of each construct was verified by Sanger sequencing.

*Pseudotyped SARS-CoV-2 variants*

The VSV-based pseudotyped SARS-CoV-2 variants were generated as described previously[2]. Briefly, HEK293T cells were transfected using 1 mg/mL of PEI with appropriate amounts of expression plasmids encoding the S protein. Following transfection, the HEK293T cells were cultured at 37°C in an atmosphere containing 5% CO2 for 24 hours. These cells were subsequently infected with VSV-G pseudotyped ΔG-luciferase (G*ΔG-luciferase, Kerafast). After incubating for two hours at 37°C, the infected HEK293T cells were washed three times and then placed in fresh medium to be cultured for another 24 hours under identical conditions. Supernatants were subsequently collected, centrifuged to remove precipitates, and aliquoted for storage at -80 °C. Prior to infection of target cells, the viral stock was incubated with 20% I1 hybridoma (anti-VSV-G) supernatant (ATCC; CRL-2700) for 1 h at 37°C to neutralize any residual VSV-G pseudotyped ΔG-luciferase.

*Pseudovirus neutralization*

Neutralization assays were performed as previously described[2,3]. Prior to each test, all pseudoviruses were titrated to equilibrate the viral input. Sera were heat-inactivated and all samples were run in triplicate in 96-well plates. Sera were four-fold serially diluted in media, starting at a 1:100 dilution. Pseudoviruses were added, and the virus-sample mixture was incubated at 37 °C for 1 hour. Control wells containing only the virus were included on all plates. Vero-E6 cells were then added at a density of $4\times10^4$ cells per well, followed by a 10-hour incubation period at 37 °C in a 5% $CO_2$ atmosphere. Afterward, the cells were lysed, and luciferase activity was measured using the Luciferase Assay System (Promega) and SoftMax Pro v.7.0.2 (Molecular Devices), following the instructions provided by both manufacturers.

*Quantification and statistical analysis*

Each assay was performed in duplicate, and the 50% neutralization titer was calculated using a five-parameter dose-response curve in GraphPad Prism v.9.2. Statistical significance between paired groups was evaluated using the Wilcoxon matched-pairs signed rank test in GraphPad Prism v.9.2. Levels of significance are denoted as follows: $*p < 0.05$, $**p < 0.01$, and $***p < 0.001$.

*Antigenic cartography*

We constructed antigenic maps from serum virus-neutralization data utilizing the technique of antigenic cartography as described previously[4]. Each antigenic map was generated using the Racmacs R package (https://acorg.github.io/Racmacs/, version 1.1.35) with 1000 optimization steps, a dilution step size of zero, and the minimum column basis parameter set to “none”. A separate antigenic map was generated for each serum timepoint. All maps were optimally aligned and overlayed with virus positions shown in Figure 1C. Separate maps for each timepoint including sera positions are shown in Figure S3.

#### Acknowledgements

This study was supported by funding from the NIH SARS-CoV-2 Assessment of Viral Evolution (SAVE) Program (Subcontract No. 0258-A709-4609 under Federal Contract No. 75N93021C00014) attributed to D.D.H., as well as the NIH, NIAID under contract number 75N93019C00051 attributed to A.G.. A.B. was supported in part by T32AI100852, Columbia Integrated Training Program in Infectious Disease Research. We express our gratitude to David Manthei, Carmen Gherasim, Victoria Blanc, Pamela Bennett-Baker, Savanna Sneeringer, Lauren Warsinske, Theresa Kowalski-Dobson, Alyssa Meyers, Zijin Chu, Hailey Kuiken, Lonnie Barnes, Ashley Eckard, Kathleen Lindsey, Dawson Davis, Aaron Rico, Gabriel Simjanovski, Mayurika Patel, and Nivea Vydiswaran of the IASO study team for supplying serum samples.

#### Author Contributions

The study was conceptualized by L.L., A.G. and D.D.H. Experiments were conducted and data analyzed by Q.W., L.L., J.H., R.Z, and A.B. Project management was handled by Q.W. Serum samples were collected by R.V., C.G., and A.G. The results were interpreted and the manuscript was written by Q.W., A.B., L.L., and D.D.H. All contributing authors have reviewed and endorsed the manuscript.

**Declaration of Interests**

D.D.H. co-founded TaiMed Biologics and RenBio, serves as a consultant for WuXi Biologics and Brii Biosciences, and is a board director at Vicarious Surgical. Aubree Gordon served as a member of the scientific advisory board for Janssen Pharmaceuticals. The remaining authors have declared no conflicts of interest

**Table S1. Summary of clinical cohorts**

| N=18 | | **No. or Mean** | **% or (range)** |
|---|---|---|---|
| Age | | 68.9 | (60, 94) |
| Female | | 8 | 44.4 |
| Male | | 10 | 55.6 |
| M1 and M2 | Pfizer | 14 | 77.8 |
| | Moderna | 3 | 16.7 |
| | AstraZeneca | 1 | 5.6 |
| M3 | Pfizer | 15 | 83.3 |
| | Moderna | 3 | 16.7 |
| M4 | Pfizer | 15 | 83.3 |
| | Moderna | 3 | 16.7 |
| B1 | Pfizer | 15 | 83.3 |
| | Moderna | 3 | 16.7 |
| B2 | Pfizer | 15 | 83.3 |
| | Moderna | 3 | 16.7 |

M1: First monovalent vaccine; M2: Second monovalent vaccine; M3: Third monovalent vaccine; M4: Fourth monovalent vaccine; B1: First bivalent vaccine; B2: Second bivalent vaccine

**Table S2. Demographics of clinical cohorts**

| Sample ID | Vaccine history | Days post M4 | Days post B1 | Days pre B2 | Days post B2 | M1-M2 Days | M2-M3 Days | M3-M4 Days | M4-B1 Days | B1-B2 Days | Age | Gender |
|---|---|---|---|---|---|---|---|---|---|---|---|---|
| L1 | BNT162b2/BNT162b2/BNT162b2/BNT162b2/Pfizer Bivalent/Pfizer Bivalent | 27 | 24 | 4 | 33 | 22 | 278 | 168 | 169 | 211 | 66 | Female |
| L2 | BNT162b2/BNT162b2/BNT162b2/BNT162b2/Pfizer Bivalent/Pfizer Bivalent | 56 | 89 | 60 | 31 | 21 | 267 | 226 | 113 | 238 | 68 | Male |
| L3 | BNT162b2/BNT162b2/BNT162b2/BNT162b2/Pfizer Bivalent/Pfizer Bivalent | 29 | 27 | 54 | 28 | 21 | 216 | 169 | 164 | 241 | 74 | Male |
| L4 | BNT162b2/BNT162b2/BNT162b2/BNT162b2/Pfizer Bivalent/Pfizer Bivalent | 20 | 64 | 15 | 32 | 21 | 244 | 192 | 160 | 225 | 64 | Female |
| L5 | BNT162b2/BNT162b2/BNT162b2/BNT162b2/Pfizer Bivalent/Pfizer Bivalent | 18 | 28 | 27 | 33 | 21 | 245 | 200 | 168 | 214 | 64 | Female |
| L6 | BNT162b2/BNT162b2/BNT162b2/BNT162b2/Pfizer Bivalent/Pfizer Bivalent | 20 | - | 43 | 34 | 21 | 231 | 191 | 162 | 233 | 65 | Female |
| L7 | BNT162b2/BNT162b2/BNT162b2/BNT162b2/Pfizer Bivalent/Pfizer Bivalent | 15 | 23 | 18 | 29 | 21 | 222 | 188 | 187 | 217 | 70 | Male |
| L8 | mRNA-1273/mRNA-1273/mRNA-1273/mRNA-1273/Moderna Bivalent/Pfizer Bivalent | 29 | 60 | 29 | 33 | 25 | 241 | 224 | 91 | 210 | 63 | Male |
| L9 | BNT162b2/BNT162b2/BNT162b2/BNT162b2/Pfizer Bivalent/Pfizer Bivalent | 75 | 18 | 25 | 33 | 21 | 264 | 216 | 123 | 220 | 63 | Male |
| L10 | AstraZeneca/AstraZeneca/BNT162b2/BNT162b2/Pfizer Bivalent/Pfizer Bivalent | 18 | 16 | 20 | 43 | 28 | 333 | 147 | 183 | 206 | 73 | Male |
| L11 | BNT162b2/BNT162b2/BNT162b2/BNT162b2/Pfizer Bivalent/Pfizer Bivalent | 43 | 22 | 5 | 34 | 21 | 252 | 203 | 182 | 181 | 66 | Male |
| L12 | BNT162b2/BNT162b2/BNT162b2/BNT162b2/Pfizer Bivalent/Pfizer Bivalent | 80 | 104 | 22 | 26 | 21 | 242 | 185 | 163 | 233 | 78 | Male |
| L13 | BNT162b2/BNT162b2/BNT162b2/BNT162b2/Pfizer Bivalent/Pfizer Bivalent | 32 | 45 | 15 | 33 | 22 | 217 | 235 | 154 | 235 | 68 | Female |
| L14 | BNT162b2/BNT162b2/BNT162b2/BNT162b2/Pfizer Bivalent/Moderna Bivalent | 21 | 57 | 58 | 32 | 21 | 230 | 178 | 164 | 225 | 66 | Female |
| L15 | BNT162b2/BNT162b2/BNT162b2/BNT162b2/Pfizer Bivalent/Pfizer Bivalent | 21 | 42 | 63 | 31 | 21 | 213 | 206 | 175 | 229 | 74 | Male |
| L16 | BNT162b2/BNT162b2/BNT162b2/BNT162b2/Pfizer Bivalent/Pfizer Bivalent | 33 | 27 | 19 | 46 | 28 | 240 | 183 | 159 | 234 | 94 | Male |
| L17 | BNT162b2/BNT162b2/BNT162b2/BNT162b2/Moderna Bivalent/Pfizer Bivalent | 15 | 21 | 5 | 21 | 21 | 141 | 154 | 234 | 222 | 64 | Female |
| L18 | mRNA-1273/mRNA-1273/mRNA-1273/mRNA-1273/Moderna Bivalent/Moderna Bivalent | 35 | 53 | 51 | 42 | 28 | 210 | 240 | 77 | 223 | 60 | Female |

M1: First monovalent vaccine; M2: Second monovalent vaccine; M3: Third monovalent vaccine; M4: Fourth monovalent vaccine; B1: First bivalent vaccine; B2: Second bivalent vaccine. One missing sample is indicated by “-”.

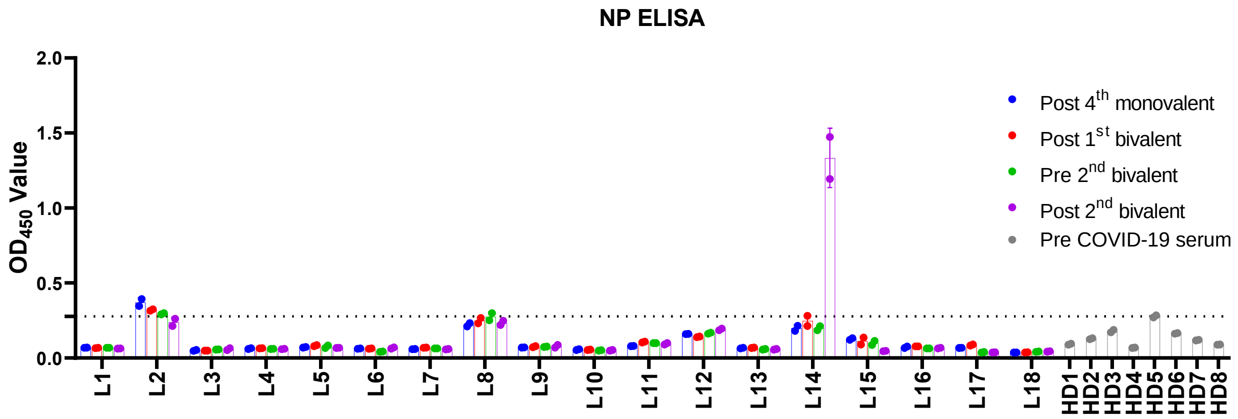

**Figure S1. Nucleocapsid binding ELISA**

Each serum timepoint from 18 individuals (L1-L18) was tested by ELISA for the presence of antibodies to the SARS-CoV-2 nucleocapsid protein (NP), indicative of prior infection. A panel of sera from healthy donors (HD1-HD8) obtained prior to the COVID-19 pandemic were used as negative controls.

#### Supplementary References

1. Simon, V. *et al.* PARIS and SPARTA: Finding the Achilles' Heel of SARS-CoV-2. *mSphere* **7**, (2022).
2. Liu, L. *et al.* Potent neutralizing antibodies against multiple epitopes on SARS-CoV-2 spike. *Nature* **584**, 450–456 (2020).
3. Wang, Q. *et al.* Alarming antibody evasion properties of rising SARS-CoV-2 BQ and XBB subvariants. *Cell* **186**, 279 (2023).
4. Wilks, S. H. *et al.* Mapping SARS-CoV-2 antigenic relationships and serological responses. *bioRxiv* 2022.01.28.477987 (2022) doi:10.1101/2022.01.28.477987.
